# Bacterially produced small molecules stimulate diatom growth

**DOI:** 10.1101/2020.11.02.365239

**Authors:** John Sittmann, Munhyung Bae, Emily Mevers, Muzi Li, Andrew Quinn, Ganesh Sriram, Jon Clardy, Zhongchi Liu

**Affiliations:** Dept. of Cell Biology and Molecular Genetics, University of Maryland, College Park, MD 20742; Dept. of Biological Chemistry and Molecular Pharmacology, Harvard Medical School, Boston, MA 02115; Dept. of Chemical and Biomolecular Engineering, University of Maryland, College Park, MD 20742

## Abstract

Diatoms are photosynthetic microalgae that fix a significant fraction of the world’s carbon. Because of their photosynthetic efficiency and high-lipid content, diatoms are priority candidates for biofuel production. Here, we report that sporulating *Bacillus thuringiensis* when in co-culture with a marine diatom *Phaeodactylum tricornutum* significantly increases the diatom cell count. Bioassay-guided purification led to the identification of two diketopiperazines (DKPs) that both stimulate *P. tricornutum* growth and increase its lipid content. RNA-seq analysis revealed upregulation of a small set of *P. tricornutum* genes involved in iron starvation response and nutrient recycling when DKP was added to the diatom culture. This work demonstrates that two DKPs produced by a bacterium could positively impact *P. tricornutum* growth and lipid content, offering new approaches to enhance *P. tricornutum*-based biofuel production. As increasing numbers of DKPs are isolated from marine microbes, the work gives potential clues to bacterially produced growth factors for marine microalgae.

**One sentence summary:** Two diketopiperazines (DKPs) produced by sporulating bacterium *Bacillus thuringiensis* stimulate diatom *P. tricornutum* growth and increase diatom lipid content.

## Introduction

Diatoms are unicellular photosynthetic algae that perform critical environmental functions with global implications. Thriving in various aquatic environments and equipped with efficient carbon concentration mechanisms, diatoms are among the planet’s most productive microalgae and fix a fifth of global carbon (Amin et al., 2012; Hildebrand et al., 2012). Diatoms fix carbon in the form of lipid droplets, a particularly useful form of biofuels, tolerate harsh environments, and perform well in large scale cultures (Hildebrand et al., 2012). Because of these features, diatoms are considered some of the most promising microalgae for biofuel production. Even with these desirable features, a scalable commercially viable microalga-derived biofuel has not yet been realized, and a better understanding of diatom physiology could increase the feasibility of diatom-derived biofuels (Hu et al., 2008; Hildebrand et al., 2012).

Diatoms and marine microorganisms have co-inhabited the oceans for more than 200 million years. The close association and interaction among them are evident by substantial horizontal gene transfer; an estimated 784 genes of bacterial origin are found in the genome of the model diatom *Phaeodactylum tricornutum* (Bowler et al., 2008). Beneficial diatom-bacterium interactions through nutrient exchange are well documented where bacteria provide the diatom with micronutrients, such as vitamin B12 or siderophores for iron uptake in exchange for dissolved organic matter produced by the diatoms (Croft et al., 2005; Boyd and Ellwood, 2010; Foster et al., 2011; Seyedsayamdost et al., 2011; Segev et al., 2016). In addition, natural compounds of marine origin are being explored to probe into diatom physiology and increase lipid production. Two small molecules penicillide and verrucarin J isolated from fungi of marine origin were recently shown to promote neutral lipid accumulation in *P. tricornutum* but unfortunately inhibited diatom growth (Yu et al., 2020). Increased knowledge on bacterium-diatom interaction may offer new strategies to enhance diatom growth and diatom-derived lipid production.

In this study, we investigated a bacterium-diatom co-culture, which showed increased *P. tricornutum* growth in the presence of *Bacillus* sp. To follow up, we co-cultured the *P. tricornutum* with a variety of Gram-positive bacteria and then counted *P. tricornutum* cells daily. Co-culturing experiments with one particular Gram-positive bacterial species, *Bacillus thuringiensis*, revealed that sporulation-mediated cell lysis of this bacterium released a small molecule metabolite capable of stimulating *P. tricornutum* cell proliferation by nearly three-folds. Purification of *B. thuringiensis* mother cell lysate and characterization of the factor led to the identification of two diketopiperazines (DKPs), the smallest possible cyclic peptides made up of only two amino acids, as being responsible for the stimulating activity. After identifying the two DKPs, we performed RNA-seq experiment of the DKP-stimulated *P. tricornutum* culture to identify cellular pathways modulated by the DKPs. This approach revealed upregulation of a small set of genes involved in iron starvation response, nutrient recycling, and high energy compound synthesis. Taken together these studies identified two bacterium-derived small molecules – members of a much larger family – that possess activities on the growth of marine diatom *P. tricornutum*, offering new approaches for enhancing algae-based biofuel production and potential clues to bacterially produced growth factors for marine microalgae.

## Results

### Discovery of bacterial-derived factor(s) that induce the growth of *P. tricornutum*

To investigate the effect of various Gram-positive bacteria on *P. tricornutum* growth, we counted *P. tricornutum* cell numbers daily in each bacterium-diatom co-culture and discovered that *Bacillus thuringiensis* (sp. *israelensis*), a member of the *Bacillus cereus* group, produced a particularly robust signal: a two to three-fold increase in *P. tricornutum* cells at stationary phase (Figure 1A, C). This ability was shown to be selective to *B. cereus* group bacteria (Figure 1A, B). Both *B. thuringiensis* sp. 4A4, and *B. thuringiensis* sp. 4Q7 also stimulated growth, but co-culturing with *B. subtilis, B. amyloliquefaciens*, or *Staphylococcus aureus* had no effect on the growth of *P. tricornutum* (Figure 1A, B). Microscopic examination of the *B. thuringiensis-* stimulated co-culture with *P*. *tricornutum* revealed the presence of *B. thuringiensis* endospores (Figure 1D, arrow), which are characteristically produced in nutrient poor L1 medium.

**Figure 1.**
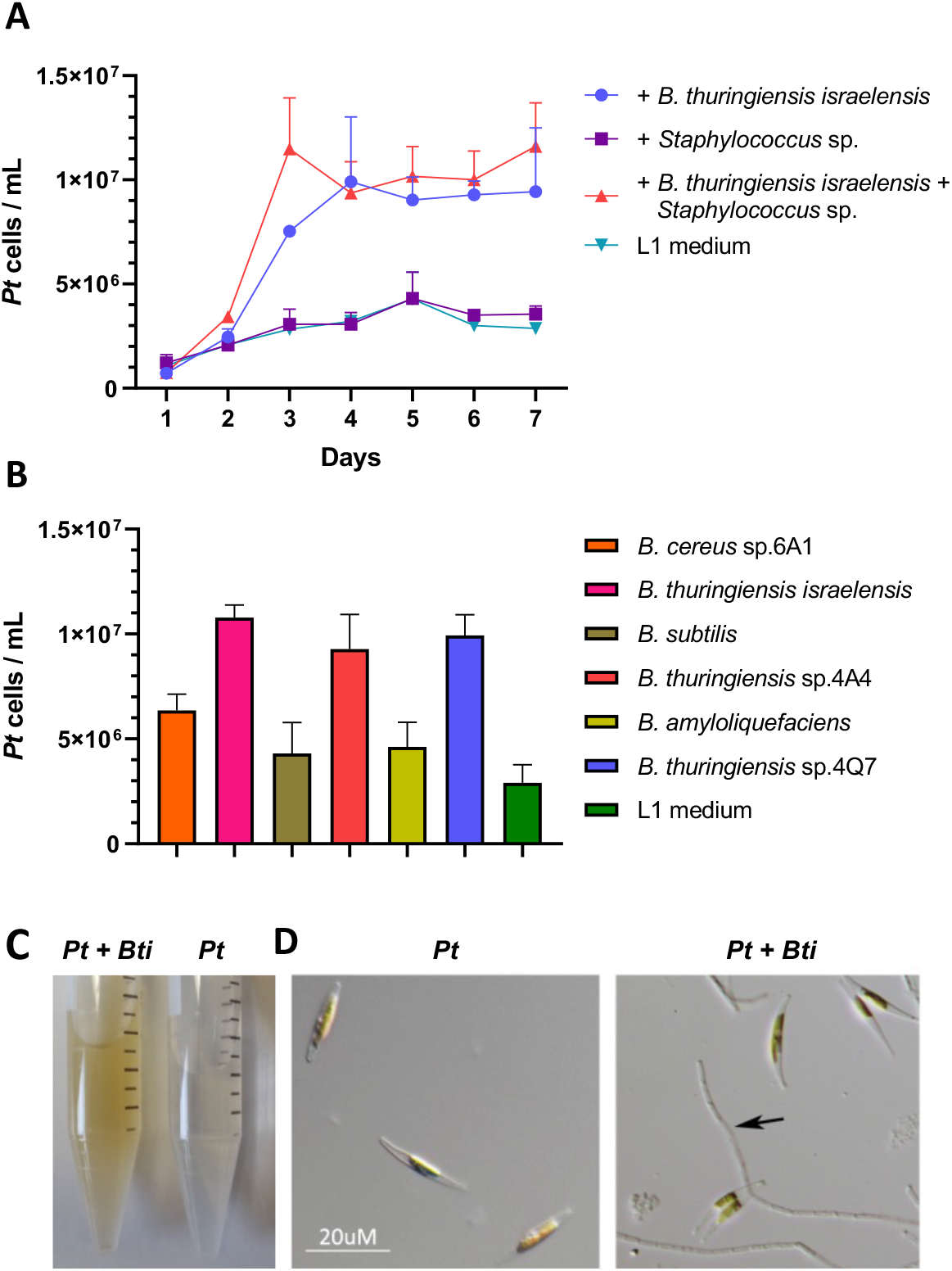
*Bacillus cereus* clade of bacteria stimulate *P. tricornutum* growth during co-culture. **A.** Growth curve of *P. tricornutum* when co-cultured with *Staphylococcus, Bacillus thuringiensis* (sp*. israelensis*), both bacteria combined, or in L1 medium (axenic condition). The bacteria are added to the *P. tricornutum* culture at day 1. The *P. tricornutum* (*Pt*) cell number (Y-axis) was quantified at each day (X-axis). **B.** *Bacillus cereus* clade members, *B. cereus* sp. 6A1 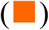, *B. thuringiensis israelensis* 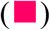, *B. thuringiensis* sp. 4A4 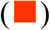, and *B. thuringiensis* sp. 4Q7 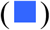 as well as *Bacillus subtilis* clade members, *B. subtilis* 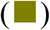 and *B*. *amyloliquefaciens* 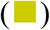 are tested for their ability to stimulate *P. tricornutum* growth. *P. tricornutum* cells (Y-axis) were counted at day 6 of co-culture. Each culture condition has 3 biological replicates. Error bars are ± 1 SD. **C**. A tube containing co-culture of *Pt* and *B. thuringiensis israelensis* (*Bti*) and a tube with *Pt* in axenic condition. The green color tube indicates higher *Pt* cell density. **D**. Microscopic images of *Pt* cells (left) and *Pt* cells in co-culture (right). A chain of *Bti* spores is indicated by an arrow. Scale bar is the same in both images.

Comparison of *P*. *tricornutum* cells and the *B*. *thuringiensis* spores over a 5-day period during the co-culture revealed that both *P*. *tricornutum* cells and *B*. *thuringiensis* spores increased rapidly between day 2-4 (Figure 2A), suggesting that sporulation directly correlates to the growth of *P. tricornutum.* In *Bacillus*, sporulation is accompanied by mother-cell lysis (Higgins and Dworkin, 2012), which would release growth stimulating intracellular factors into the culture media. To test this hypothesis*, P. tricornutum* was treated with *B. thuringiensis*’s spores or the mother cell lysate, respectively, when the spores and cell debris were removed. Growth stimulation was only observed when using the mother cell lysate (Figure 2B). Mechanically lysed *B. thuringiensis* cells growing in nutrient rich LB medium (no sporulation) by sonication and subsequent filtration was unable to stimulate diatom growth (Figure 2C), supporting that sporulation under nutrient deplete conditions is required for the production of the stimulating factors. The mother cell lysate was heat-treated by autoclaving (121 °C, 30 min), which resulted in loss of stimulating activity (Figure 2D). Pretreatment of the *B. thuringiensis* lysate with Proteinase K did not affect the stimulating effect of the lysate (Figure S1). Thus, a small molecule is likely responsible for the growth promoting effect.

**Figure 2.**
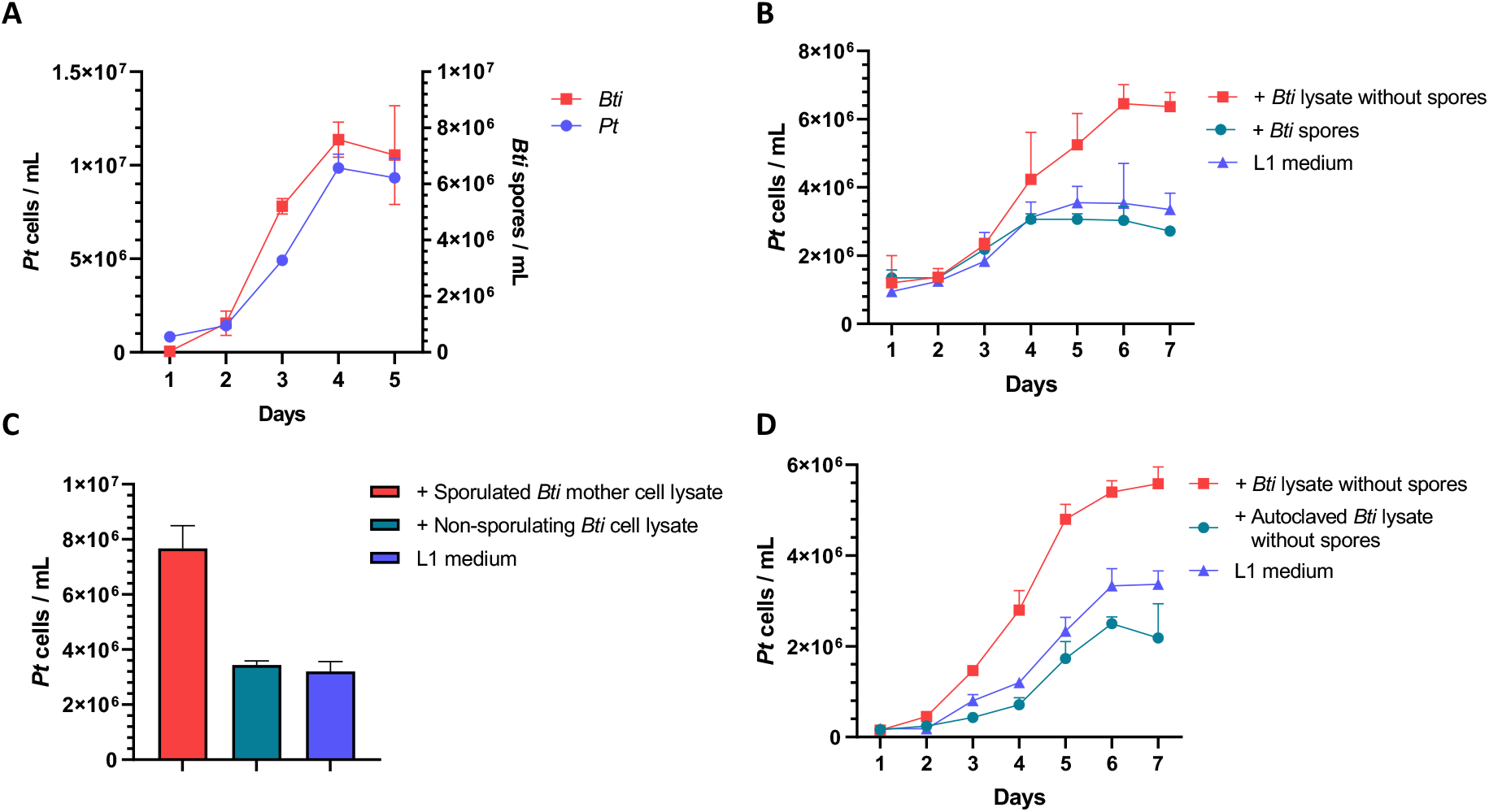
The mother-cell lysate of *B. thuringiensis israelensis* contains a heat labile stimulator. **A.** *P. tricornutum* 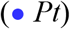 cell counts (left Y-axis) and *B. thuringiensis israelensis* 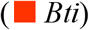 spore counts (right Y-axis) in co-cultures, showing correlation between diatom cells and bacterial spores. **B.** *P. tricornutum* cell counts in culture with *B. thuringiensis israelensis* (*Bti*) mother cell lysate (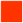, spores removed), the spores 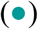, and L1 medium 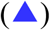, respectively. **C.** *P tricornutum* cell counts in cultures containing sporulated *B. thuringiensis israelensis* (*Bti*) mother cell lysate 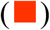, non-sporulating *B. thuringiensis israelensis (Bti)* cell lysate 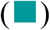, and the axenic control in L1 medium 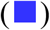. Cell counts were done at day 7 of co-culture. **D.** *P. tricornutum* cell counts in culture with *B. thuringiensis israelensis (Bti)* mother cell lysate 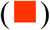, autoclaved *B. thuringiensis* mother cell lysate 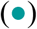, or the axenic control in L1 medium 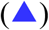. Each culturing condition contained 3 replicates. Error bars are ± 1 SD.

### Identification of the growth stimulating factor(s)

The growth stimulating factor was captured from the *B. thuringiensis* mother cell lysate using a mixture of XAD resins, and the crude extract was fractionated into five reduced complexity fractions using reverse-phase solid phase extraction (RP-SPE) columns. Growth-stimulating activity was in the water fraction, indicating that the active small molecules are highly polar. Further fractionation using RP high performance liquid chromatography (HPLC), yielded twenty-five additional fractions, which were screened for growth stimulation of *P. tricornutum.* Only two fractions exhibited activity (Figure 3A, B). Comprehensive analysis of UV, mass, and NMR data revealed that the active fractions contained two cyclic dipeptides (**1** and **2**) belonging to diketopiperazine (Figure 3C, D).

**Figure 3.**
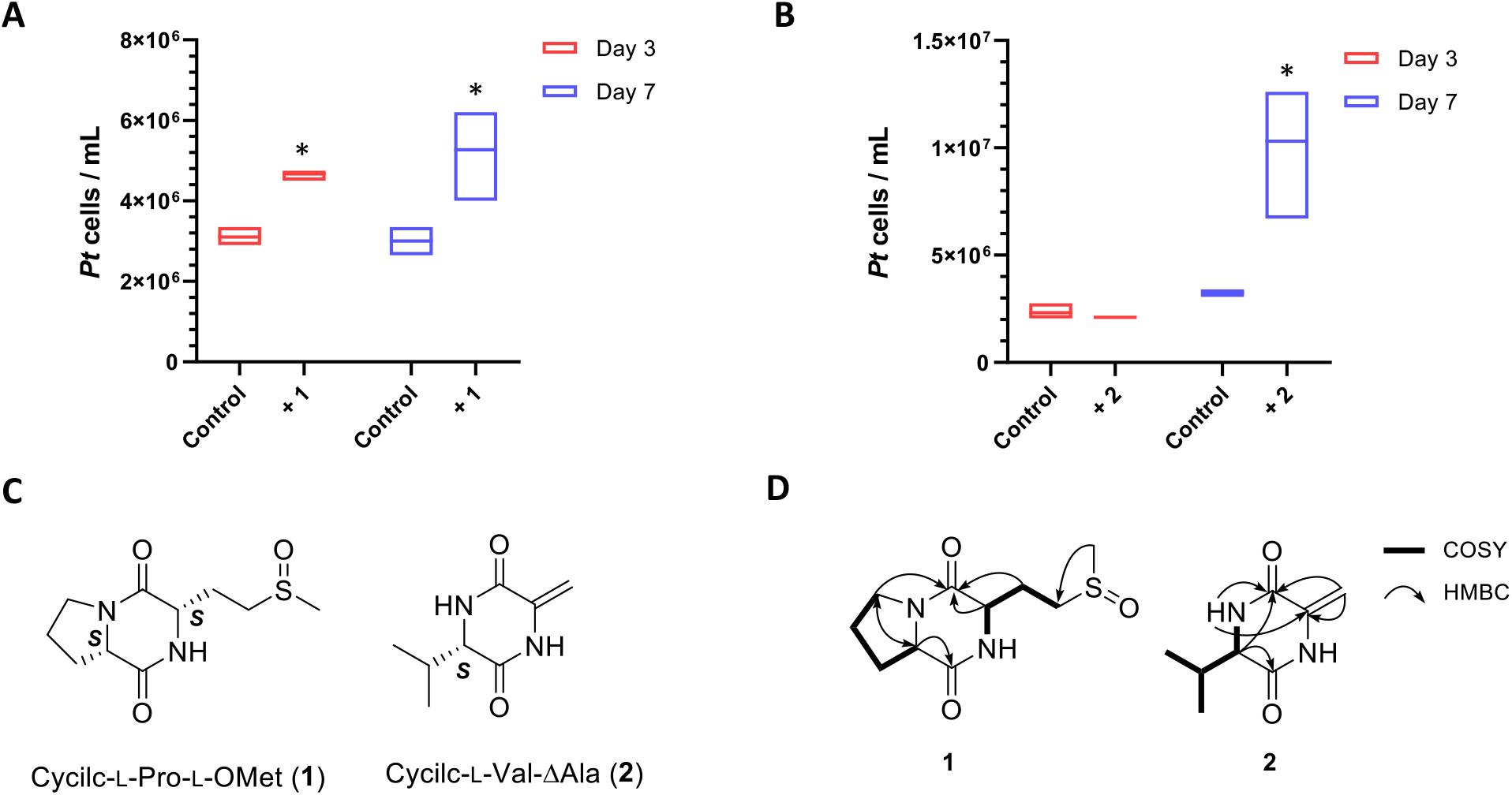
Identification of two diketopiperazines as the growth simulators from *B*. *thuringiensis.* **A**. The growth enhancing effect of **1** was significant at day 3, and day 7 cultures. **B**. Significant stimulating effect of **2** was shown at day 7 culture. Each culturing condition has 3 biological replicates. Error bars are ± 1 SD. “*” indicates P-value <0.05. **C**. Structures of cyclic-l-Pro-l-*O*Met (**1**) and cyclic-l-Val-ΔAla (**2**). **D.** Key COSY and HMBC correlations of **1** and **2**.

Compound **1** was purified as a white powder having a molecular formula of C10H16N2O3S based on HR-ESIMS (obsd. [M+H]^+^ *m/z* 245.0972, Δ 2.8 ppm), a formula that requires four degrees of unsaturation. Interpretation of the 1D and 2D NMR data (^1^H, ^13^C, HSQC, COSY, and HMBC) (Table S1) of **1** revealed two carboxyl carbons (*δ*_C_ 175.2 and 168.8), two α-carbons (*δ*_C_ 61.7 and 56.4), five aliphatic carbons (*δ*_C_ 50.2, 48.1, 30.4, 25.2, and 24.6), and one singlet methyl (*δ*_C_ 39.2) make up three distinct spin systems. The first spin system – with COSY correlations between H-3 (*δ*_H_ 3.57), H-4 (*δ*_H_ 2.08/1.98), H-5 (*δ*_H_ 2.36/1.97), and *α*-proton H-6 (*δ*_H_ 4.36) – is indicative of a proline residue. The second spin system contained COSY correlations between an α-methine H-9 (*δ*_H_ 4.53) and two methylenes, H-10 (*δ*_H_ 2.38/2.31) and H-11 (*δ*_H_ 2.97). A key HMBC correlation from the singlet methyl group, H-13 (*δ*_H_ 2.74), to C-11 led to the assignment of the second partial structure as methionine sulfoxide. Additional HMBC correlations from H-9 to C-1 (*δ*_C_ 168.8) and from H-6 to C-7 (*δ*_C_ 175.2) connected the two partial structures and established the planar structure of **1** as cyclic-l-Pro-l-*O*Met, a diketopiperazine (DKP) (Figure 3C, D).

Compound **2** has a molecular formula of C_8_H_13_N_2_O on the basis of HR-ESIMS (obsd. [M+H]^+^ *m/z* 169.0981, Δ 1.2 ppm) and based on the 1D NMR data **2** appeared to be another DKP (Table S2). Analysis of the chemical shifts and UV spectrum indicated **2** incorporated a terminal methylene (*δ*_C_/*δ*_H_ 98.9/5.17 and 4.76) and geminal dimethyl moiety. Detailed analysis of the 2D NMR dataset revealed the planar structure of **2** is cyclic-l-Val-ΔAla (Figure 3C, D).

The absolute stereoconfigurations of the three chiral amino residues in **1** and **2** were determined by Marfey’s analysis. Each DKP was hydrolyzed and then derivatized with l-FDAA (1-fluoro-2-4-dinitrophenyl-5-l-alanine amide) (Szókán et al., 1988). Comparison of the reaction mixture to authentic standards using a HR-LCMS revealed that all amino acids correspond to l-amino acids (Table S3).

Due to low production yields of both **1** and **2** by the producing bacterium and a desire to rule out an active impurity, we sought an alternative source for each metabolite in order to obtain enough material for unambiguous biological evaluation. Compound **2** was commercially available (BocSciences), but we had to synthesize **1**. The synthesis involved two-steps and traditional peptide coupling chemistry (Campbell and Blackwell, 2009). Both metabolites stimulated diatom growth in dose-dependent manners with minimal active concentrations of 100 nM and 35 nM for compounds **1** and **2**, respectively (Figure 4A). Importantly, addition of individual amino acids that constitute the DKP’s building blocks, even at high concentrations (0.7 - 10 mM), had no effect on the diatom growth (Figure 4B). Furthermore, chemical profiling revealed that all growth stimulatory *B. thuringiensis* strains produced between 4 to 20 μM of both **1** and **2** in spent media, with *B. thuringiensis* sp. 407 producing the highest amounts of each of the DKPs (Figure 4C, D).

**Figure 4.**
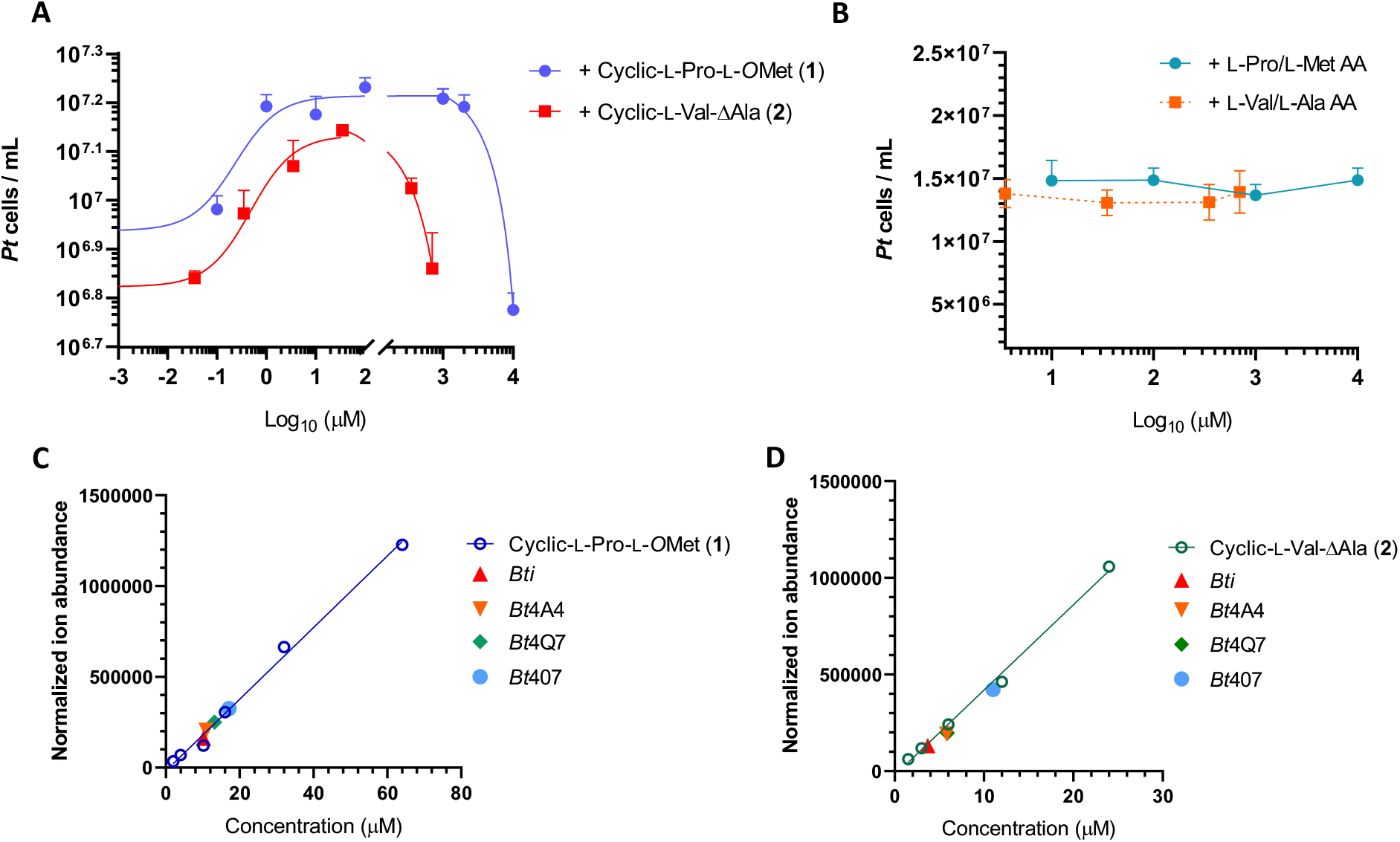
The bacteria-derived DKPs (cyclic-l-Pro-l-*O*Met and cyclic-l-Val-ΔAla) stimulate *P. tricornutum* growth in a dose-dependent manner. **A.** *P. tricornutum* cell numbers when growing in culture containing different concentrations of cyclic-l-Pro-l-*O*Met (**1**) and cyclic-l-Val-ΔAla (**2**). Cell counts were done at day 7, when effective stimulation was established for both DKPs. **B.** Solutions containing mixed pair of single l-amino acids (l-Pro/l-Met AA and l-Val/l-Ala AA) failed to stimulate *P. tricornutum* growth at different concentrations. Cell counts were done at day 8 of culture. Each culturing condition has 3 biological replicates. Error bars are ± 1 SD. **C.** Absolute quantification of cyclic-l-Pro-l-*O*Met (**1**) and **D.** Absolute quantification of cyclic-l-Val-ΔAla (**2**) in *B*. *thuringiensis israelensis* 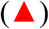, *B*. *thuringiensis* sp.4A4 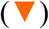, *B*. *thuringiensis* sp.4Q7 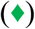, *B*. *thuringiensis* sp.407 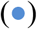. The calibration curve is also shown.

### Transcriptome profiling of DKP-stimulated *P. tricornutum* cultures

To gain insights into the mechanism of DKP-induced *P. tricornutum* growth, we examined gene expression changes in *P. tricornutum* using RNA-seq analysis. *P. tricornutum* grown in L1 media with and without the addition of l-Pro-l-*O*Met (**1**) (100 μM) were harvested on day 3 and day 6. Day 3 is when the two cultures are about to deviate in growth rate (Figure S2A), which allows detection of earliest gene expression changes. Day 6 is when both cultures are at the exponential phase (Figure S2A), thus minimizing differences of the two cultures caused by differential health status. Surprisingly, principal component analysis (PCA) of the RNA-seq data revealed that while the DKP treated samples are distinct from untreated controls at day 3, DKP-treated samples are highly similar to untreated controls at day 6 (Figure S2B), suggesting that the effect of the DKP is transient and the DKP-treated and untreated cultures are otherwise highly similar.

Thirty-nine differentially expressed genes (DEGs) were identified between DKP-treated and untreated samples at day 3 (Data S1) with 32 up-regulated and 7 down-regulated genes. Consistent with PCA analysis, no DEG was identified at day 6 between the two conditions. Among the 32 up-regulated DEGs at day 3, twelve have annotation of putative function (Figure 5A); most were previously shown to be induced under iron starvation conditions in diatoms (Allen et al., 2008; Lommer et al., 2012), including ISIP1, ISIP2A, ISIP2B, ISIP3, and a Cell Surface protein with 25% identity to ISIP2B. In addition, two induced ferric reductase genes (FREs) may function to reduce chelated-Fe^3+^ to Fe^2+^ (Allen et al., 2008; Kazamia et al., 2018; Coale et al., 2019); Fe^2+^ is more soluble and available to cells. It maybe that the up-regulation of these genes helps *P. tricornutum* cells survive and perhaps even thrive under iron-limiting conditions.

**Figure 5.**
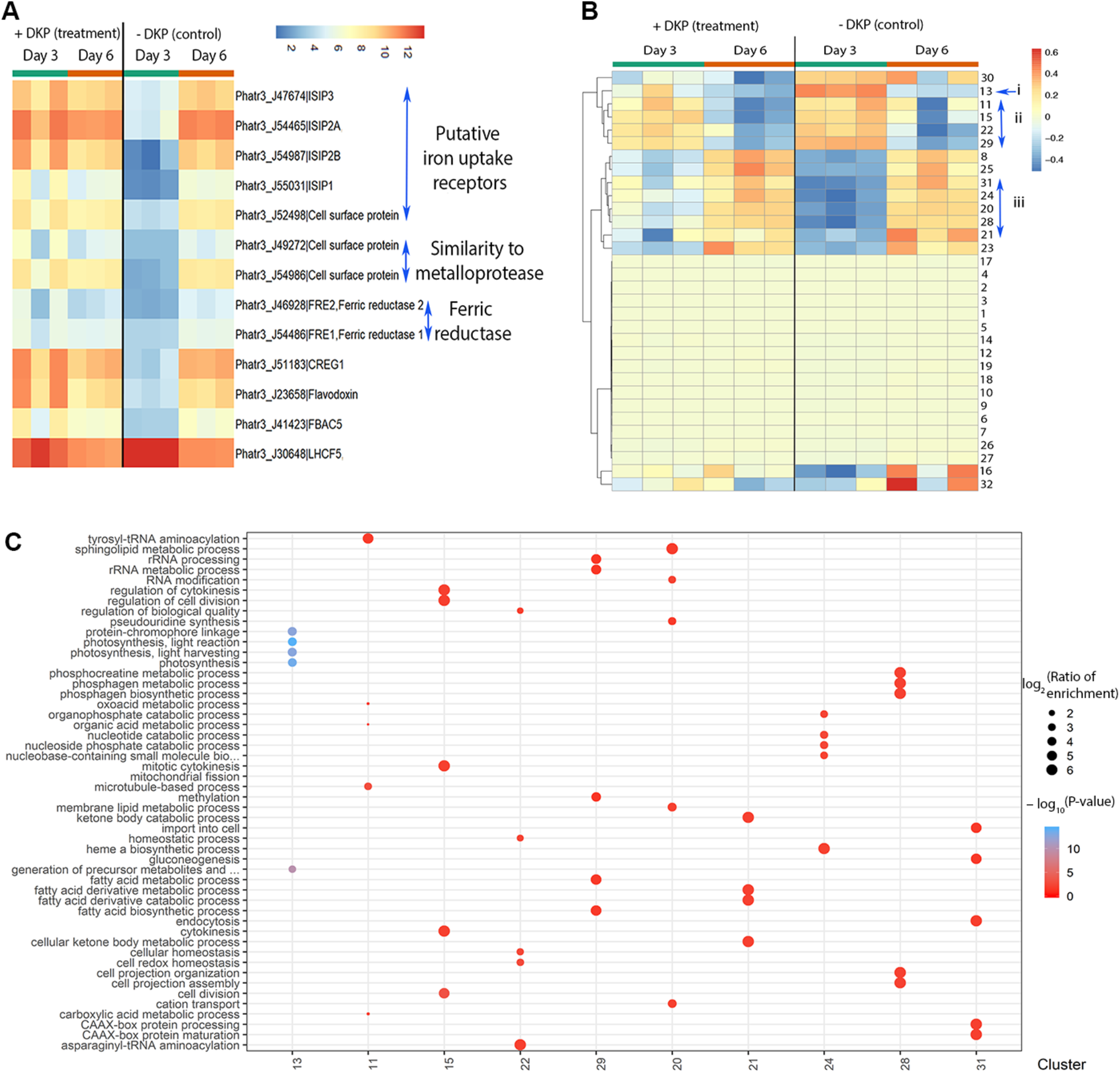
Transcriptome analyses revealed reduced expression of photosynthesis genes and increased expression of iron uptake genes in DKP-treated *P. tricornutum* at day 3. **A.** Heatmap of differentially expressed genes between DKP-treated and not treated samples at day 3. Each row is a differentially expressed gene. Each column represents one of the three biological replicates. Scale bar represents relative gene expression (log2 (tpm+1). **B**. Heatmap of cluster eigen genes. Each row represents one of the 32 co-expression clusters. Each column represents eigen gene value for each of the three biological replicates. Scale bar: cluster eigengene expression value which equals to the first principle component of the gene expression value in the cluster. **C.** Enriched GO terms belonging to clusters in group i, ii and iii shown in **B**.

In addition to differential gene expression, a co-expression network was constructed using Weighted Gene Co-expression Network Analysis (WGCNA) (Langfelder and Horvath, 2008) and enriched Gene Ontology (GO) terms were obtained for each cluster (Data S2). Several co-expression clusters, 11, 15, 22, and 29 (group ii; Figure 5B) are activated at day 3 in treated and untreated samples alike; their enriched GO terms include cell division, metabolic processes, cellular homeostasis, and fatty acid metabolism, indicative of healthy cell growth during exponential phase. In contrast, cluster 13 (group i; Figure 5B), downregulated at day 6 under both conditions, is precociously downregulated at day 3 in cyclic-l-Pro-l-*O*Met-treated sample. Enriched GO term for cluster 13 is photosynthesis (Figure 5C), indicating that DKP-treated *P. tricornutum* cells may have reduced photosynthesis at day 3.

Most significant difference between DKP-treated and untreated samples are shown by group iii clusters, 31, 24, 20, 28, and 21 (Figure 5B, C); they show upregulation at day 3 in DKP-treated samples. Cluster 20 is enriched in cation transport, cluster 21 is enriched in mitochondrial fission, cellular ketone body metabolism, and fatty acid derived catabolism, suggesting a metabolic shift to the mitochondrial system (a phenomenon also observed under iron limiting conditions) (Allen et al., 2008). Cluster 24 and 28 are enriched in GOs related to nutrient recycling (nucleotide metabolic process), synthesis and breakdown of high energy storage compounds (phosphagen metabolic process and phosphocreatine metabolic process), and gluconeogenesis, a pathway for glucose synthesis from non-carbohydrate carbon substrates.

Combined, the data suggest that the DKPs, likely produced upon sporulation, trigger stress responses in *P. tricornutum.* Bacterial stress factors induce diatom stress responses. That the expression of DKP-treated samples at day 3 is similar to day 6 of both DKP-treated and untreated control could indicate that DKP may precociously trigger a stress response normally activated when nutrients are being exhausted.

### DKP changes composition and yield of fatty acids in *P. tricornutum*

While the *B. thuringiensis* lysate can effectively and inexpensively increase *P. tricornutum* growth, it is not known if such treatment would negatively impact lipid production in *P. tricornutum.* This is important as diatoms are regarded as one of the most promising sources of biofuel. To address this question, the lipid composition and yield were analyzed using gas chromatography mass spectrometry (GCMS) to identify and quantify the fatty acids in cultures stimulated with the *B. thuringiensis* cell-free lysate. At day 7, *P. tricornutum* cells were centrifuged and the pellet was subjected to extraction. The result revealed increased levels of three key components of biodiesel - palmitoleic acid (C16:1), oleic acid (C18:1) and linoleic acid (C18:2) - in *B. thuringiensis* lysate-stimulated *P. tricornutum* (Figure 6A) as well as an increase in beneficial dietary lipids, eicosapentaenoic acid (EPA) (C20:5) and palmitoleic acid (C16:1) (Figure 6A). Interestingly, all the unsaturated fatty acids increase in relative and absolute abundance, whereas the saturated fatty acids do not (Figure 6A). Shifting to unsaturated fatty acid production and enhanced overall lipid productivity have been reported in *P. tricornutum* as a stress response upon cold temperature or nitrogen limitation (Jiang and Gao, 2004; Zulu et al., 2018).

**Figure 6.**
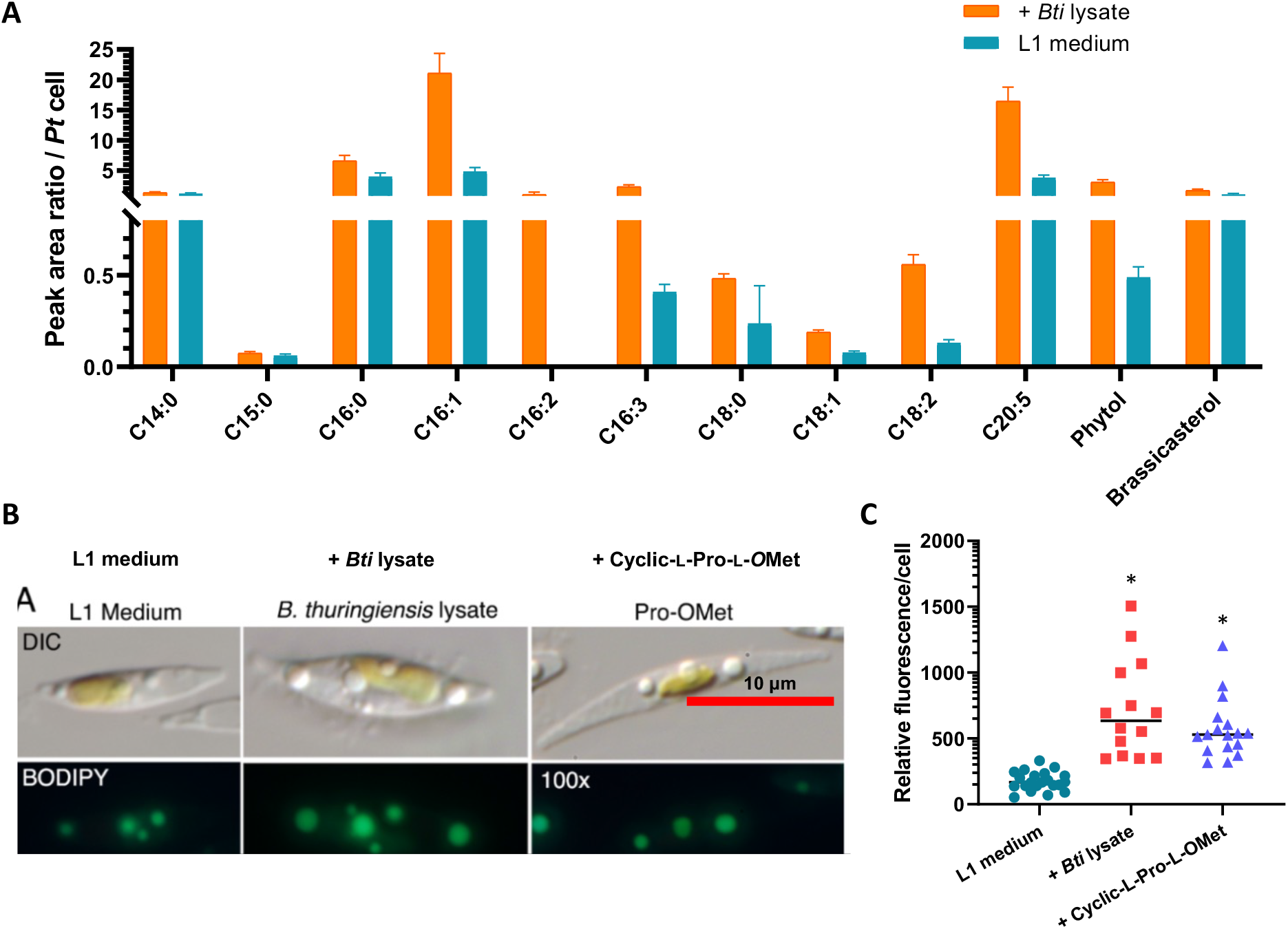
Stimulated *P. tricornutum* cells have higher lipid content and higher yield of beneficial fatty acids. **a**. Fatty acid fractions of seven-day *P. tricornutum* culture grown in L1 medium or L1 medium containing *B. thuringiensis israelensis* lysate measured by GC-Mass Spectrometry. Values represent normalized fatty acid production per cell of each culture as peak area ratio versus the internal standard. Cells grown in L1 media averaged 4.35E+06 *P. tricornutum* cells per mL, while cells grown in L1 + *B. thuringiensis israelensis* lysate averaged 1.34E+07 cells per mL. **b**. Differential Interference Contrast (DIC) microscopic images (top panel) and fluorescent microscopic images (bottom panel) of BODIPY-stained *P. tricornutum* cells at stational phase (12 days). **c**. Relative Fluorescence/cell (Y-axis) is equal to Integrated Density (see Methods) based on BODIPY staining of *P. tricornutum* cells at stationary phase. Black horizontal bars represent the average Integrated Density of cells in the experimental condition. N= 23 (*L1* medium), 30 cells (L1 medium + *B. thuringiensis israelensis* lysate), and 47 (L1 medium + cyclic-l-Pro-l-*O*Met).

The effect of the DKP on *P. tricornutum* lipid composition was also tested by staining stationary phase *P. tricornutum* cells with BODIPY 505/515, a fluorescent stains for neutral lipids (Wu et al., 2014). *P. tricornutum* cells grown in L1 medium containing 100 mM cyclic-l-Pro-l-*O*Met (**1**) or cell-free lysate of *B. thuringiensis* exhibited higher relative fluorescence per cell than the *P. tricornutum* control cultured in L1 medium only (Figure 6B-C). These results suggest that the DKP and *B. thuringiensis* lysate not only stimulate *P. tricornutum* growth but also increase neutral lipid accumulation.

## Discussion

In the current study, we showed that *Bacillus cereus* group of bacteria, upon sporulation and mother cell lysis, release small molecules into the medium, which possess the ability to stimulate *P. tricortunum* growth. Bioassay-guided purification led to the identification of two diketopiperazines (DKPs), cyclic-l-Pro-l-*O*Met (**1**) and cyclic-l-Val-ΔAla (**2**), which are able to largely replicate the growth-stimulating effect on diatom at nM concentrations. DKPs contain a heterocyclic 6-membered ring, which provides metabolic stability, protease resistance, and conformational rigidity (Borthwick, 2012; Giessen and Marahiel, 2015).

DKPs have been isolated from marine-derived bacteria, fungi, mollusk, sponge and red algae (Huang et al., 2014). Recently, cyclic-l-Pro-l-*O*Met (**1**) is among 32 DKPs identified from five bacterial strains isolated from marine sediments (Harizani et al., 2020). However, the biological function of most of these naturally produced DKPs remain elusive. The only known diatom-derived DKP is a diproline that acts as a pheromone to coordinate attraction and mating in the benthic pennate diatom *Seminavis robusta* (Gillard et al., 2013). However, sexual reproduction in *P. tricornutum* is currently not well understood, any similar roles will need further investigation. In soil, a series of proline-derived DKPs produced by *Pseudomonas aeruginosa* were found to mimic the effect of plant hormone auxin and affect root development in *Arabidopsis thaliana* (Ortiz-Castro et al., 2011). This and the discovery reported here suggest that DKPs are naturally produced from soil microbes as well.

The two DKPs identified in this study exhibited dose-dependent effect on *P. tricornutum* growth (Figure 4A). At a low concentration of 100 nM for cyclic-_L_-Pro-_L_-*O*Met (**1**) or 35 nM for cyclic-l-Val-ΔAla (**2**), the DKPs lose their growth promoting effect. Half maximal effective concentrations EC50 for growth stimulation were shown to be 320 nM for cyclic-_L_-Pro-_L_-*O*Met (**1**) and 690 nM for cyclic-_L_-Val-ΔAla (**2**). When the DKP concentration exceeds 10 mM or 700 μM for cyclic-l-Pro-l-*O*Met (**1**) or cyclic-l-Val-ΔAla (**2**) respectively, growth inhibition was instead observed (Figure 4A). This result suggests that the two DKPs maybe important regulatory factors instead of just being essential nutrients.

The absolute concentration of cyclic-l-Pro-l-*O*Met (**1**) and cyclic-l-Val-ΔAla (**2**) in the *B*. *thuringiensis israelensis* lysate was quantified to be 10 μM and 4 μM for cyclic-l-Pro-l-*O*Met (**1**) and cyclic-l-Val-ΔAla (**2**) respectively (Figure 4C, D), which are within the optimal range for the growth promotion. However, it is difficult to determine if the two DKPs fully recapitulate the effect of *B. thuringiensis* lysate (or *B. thuringiensis* cells) as the microenvironment surrounding *P. tricornutum* in the culture may affect local concentration and the effectiveness of DKP action.

Although the exact cellular mechanism of the DKPs on *P. tricornutum* is unknown, our genomewide transcriptome analysis revealed that the majority of the upregulated DEGs are associated with iron-limiting conditions. Iron-limiting was associated with decreased photosynthesis in *P. tricornutum* (Greene et al., 1991), which could explain reduced photosynthesis indicated by cluster 13 eigengene expression at day 3 (Figure 5B, C). However, iron-limiting conditions are normally associated with slower diatom growth (Zhao et al., 2018), which is in contrast to higher diatom growth shown here. One possible interpretation is that DKPs induce short term maximum iron uptake and usage by inducing iron-limiting responses in *P. trocornutum* even under iron-sufficient conditions.

One of the main bottlenecks for algal-derived biofuels is algae cultivation and accumulation of enough biomass (Khan et al., 2018; Sajjadi et al., 2018). Most industrial cultivations of microalgae in axenic photobioreactors or low-cost open nutrient-poor ponds. The slow growth rate of algae in nutrient poor axenic medium, open pond, or aquatic conditions has limited its potential for sustainable biofuel production. *P. tricornutum* is a superb model organism for investigations that may lead to innovative solutions for the stated limitations. The available molecular genetic tools in *P. tricornutum* enables genetic engineering targeting genes in the lipid biosynthetic pathway to increase biomass (Daboussi et al., 2014; Serif et al., 2018; Wang et al., 2018). The results reported here provide another approach, the *B. thuringiensis* lysate can be inexpensively produced to stimulate *P. tricornutum* growth as well as increase neutral lipid production, leading to enhanced biomass accumulation without any observed negative impact on lipid content. Future work pairing our bacterial and chemical approaches with targeted genetic manipulation will allow for a greater understanding and improved benefit for biofuel production.

## Methods

### General experimental procedures

UV spectra were acquired by using a Ultrospec 5300-pro UV/Visible spectrophotometer. Optical rotations were recorded on a JASCO P-2000 polarimeter (sodium light source) with a 1 cm cell. NMR spectra were recorded by a Bruker Avance 500 MHz spectrometer. Electrospray ionization (ESI) low–resolution LC/MS data were obtained by using an Agilent Technologies 6130 quadrupole mass spectrometer coupled with an Agilent Technologies 1200–series HPLC. High– resolution ESI (HR–ESI) mass spectra were acquired on an Agilent LC-q-TOF Mass Spectrometer 6530-equipped with a 1290 uHPLC system. HPLC purifications were performed by using Agilent 1100 or 1200 series LC systems equipped with a photo-diode array detector.

### *Bacillus* strains and culture conditions

*Bacillus thuringiensis israelensis* was isolated from spores in the larvicide Gnatrol (Valent, United States). *Bacillus subtilis* sp. 168 and *Bacillus amyloliquefaciens* were from Dr. Wade Winkler at the University of Maryland, College Park. *Bacillus cereus* sp. 6A1 *, Bacillus thuringiensis* sp. 4A4, and *Bacillus thuringiensis* sp. 4Q7 were purchased from the Bacillus Genetics Stock Center (BGSC). *Bacillus thuringiensis* sp. 4D4 and *Bacillus thuringiensis* sp. 407 were from Dr. Vincent Lee. All *Bacillus* species were cultured in LB medium. A single colony of *Bacillus* was added to 10 mL LB, which was grown at 37 °C shaking at 200 rpm overnight.

### *Phaeodactylum tricornutum* strain, culture condition, and cell counts

*P. tricornutum* is cultured in the L1 medium, a salt water solution containing 32 g/L of Instant Ocean sea salt supplemented with 1mL (per L salt water) trace metal solution (NCMA) as well as 8.82 x 10-^4^ M NaNO_3_, 3.62 x 10^-5^ M NaH_2_PO_4_·H_2_O, and 1.06 x 10^-4^ M Na_2_SiO_3_·9H_2_O.

*P. tricornutum* strain CCMP632 was obtained from NCMA (Provasoli-Guillard National Center for Marine Algae and Microbiota, East Boothbay, ME). *P. tricornutum* cells were grown in non-shaking L1 medium at 22 °C with a light intensity of 50 μmol/m^2^/s until mid-exponential phase, and 100 μL of such *P. tricornutum* culture was transferred to 2 mL L1 medium in 15 mL Falcon tubes with a tapered bottom. Overnight *B. thuringiensis* culture (100 μL), or control medium, or nutrients were respectively added into the same 2 mL culture. Co-cultures were grown without agitation at 22 °C with a light intensity of 50 μmol/m^2^/s. *P. tricornutum* cells at any growth stage could stimulated by adding the stimulating agents.

All cell counts of *P. tricornutum* were performed using a Hauser Scientific Levy Hemacytometer. 7.5 μL of cell culture was loaded into the well. Cells occupying the 1 mm x 1 mm x 0.1 μL grid were counted in the experiment. The cell counts were then scaled up to indicate “number of *P. tricornutum* cells per mL.”

### *Bacillus thuringiensis* cell lysate production

To generate the *B. thuringiensis* mother cell lysate, 1.25 mL of *B. thuringiensis* overnight culture in LB medium was added to 25 mL of fresh L1 medium. The nutrient poor bacterial culture was kept at room temperature without shaking for two weeks to enable sporulation. The culture was spun down and the supernatant was filtered through a 0.22 μm filter to yield spore-free lysate. The spores in the pellet were washed three times in distilled water and resuspended in 1.25 mL of L1 medium; 100 μL was added to 2 mL of L1 medium for co-culture with 100 μL of *P. tricornutum.*

To generate mechanically lysed nonsporulating *B. thuringiensis* cells, 1.25mL of *B. thuringiensis* was removed from a 25 mL overnight culture in LB, spun down, and resuspended in 350 μL of L1 medium. The cellular suspension was sonicated using the Microson Ultrasonic Cell Disruptor (Misonix) at setting 12 for 10 x 1-sec pulses. The shredded cells were resuspended in 25 mL of L1 medium and filtered through a 0.22 μm filter. An aliquot (2 mL) of mechanically generated lysate was transferred into 15 mL Falcon tube followed by the addition of 100 μL *P. tricornutum* culture at exponential phase.

### Proteinase K treatment of mother cell lysate

To pre-treat the *Bacillus* mother cell lysate with Proteinase K, 2 μL of Proteinase K (20mg/mL) was added to 2 mL *Bacillus* mother cell lysate, which was allowed to sit for one week at 22 °C. Afterwards, 80 μL of water containing 4.1 mg of glycine was added to the *Bacillus* lysate*_*to inhibit the Proteinase K. At this point, 100 μL *P. tricornutum* culture was added to the 2 mL lysate pretreated with Proteinase K.

### Identification of DKP from *B. thuringiensis israelensis* lysate

*B. thuringiensis* strain was cultivated in 5 mL of LB medium (5 g yeast extract, 10 g peptone, and 5 g sodium chloride in 1 L distilled water) in a 15 mL falcon tube. After cultivating the strain for 3 days on a rotary shaker at 180 rpm at 30 °C, 5 mL of the *B. thuringiensis* liquid culture was inoculated to 500 mL of L1 medium [1 mL NaNO_3_ (75.0 g/L dH_2_O), 1 mL NaH_2_PO_4_·H_2_O (5.0 g/L dH_2_O), 1 mL Na_2_SiO_3_·9H_2_O (30.0 g/L dH_2_O) in 1 L artificial seawater] in a 1 L Pyrex media storage bottle. To induce sporulation and generate *B. thuringiensis* mother-cell lysate, the *B. thuringiensis* was incubated at 22 °C. The culture was allowed to sporulate for 21 days to generate the mother-cell lysate. Supernatant from 8 L culture for the *B. thuringiensis israelensis* was loaded on a column containing 100 g of XAD4 and 100 g of XAD7. After loading the supernatant, the column was washed with 500 mL of DI water and then the crude extract was eluted with 500 mL of 100% MeOH. The MeOH was evaporated to yield a crude extract. The crude extract was then resuspended in approximately 50 mL of 100% DI water and loaded on a 10 g RP C_18_ Solid Phase Extraction (SPE) cartridge and fractionated using the following solvent system: 100% H_2_O (fraction A), 20% (fraction B), 40% (fraction C), 60% (fraction D), 80% (fraction D), and 100% (fraction E) MeOH/H_2_O. The growth simulation activity was only detected in fraction A - 100% water fraction.

Fraction A (100% water fraction) was then subjected to reversed-phase prep-HPLC (Luna® Phenyl-Hexyl: 250 × 21.2 mm, 5 μm) with the following linear gradient elution: 5% acetonitrile (ACN)/95% H_2_O to 20% ACN/80% H_2_O over 40 min with a flow rate of 10 mL/min. Fractions were collected every 2 min between 5 min and 55 min, generating 25 refined fractions. Growth stimulating activity was detected in fractions A13 (31 min) and A19 (43 min) yielding cyclic-l-Pro-l-*O*Met (**1**-2.5 mg) and cyclic-l-Val-ΔAla (**2**-1.2 mg), respectively.

Cyclic -l-Pro-l-OMet (**1**). [α]_D_ −125.8 (c 0.1, MeOH); UV (MeOH) *λ_max_* (log ε) 210 (4.12) nm; HR-ESIMS *m/z* 245.0972 [M+H]^+^ (calcd. for C_10_H_17_N_2_O3S 245.0965); ^1^H NMR (500 MHz, D_2_O) 4.53 (t, *J* = 5.0, 1H), 4.36 (t, *J* = 8.0, 1H), 3.57 (m, 2H), 2.97 (m, 2H), 2.74 (s, 3H), 2.38-2.31 (m, 3H), 2.08 (m, 1H), 1.98-1.97 (m, 2H); ^13^C NMR (125 MHz, D_2_O) *δ*_C_ 175.2, 168.9, 61.8, 56.5, 50.3, 48.1, 39.3, 30.5, 25.3, 24.6.

Cyclic -l-Val-ΔAla (**2**). [α]_D_ −113.2 (c 0.1, MeOH); UV (MeOH) *λ_max_* (log ε) 220 (4.08) nm, 240 (3.96); HRESIMS *m/z* 169.0981 [M+H]^+^ (calcd for C_8_H_13_N_2_O_2_ 169.0983); ^1^H NMR (500 MHz, DMSO-*d*β) *δ*_H_ 10.52 (brs, 1H), 8.36 (s, 1H), 5.17 (s, 1H), 4.77 (s, 1H), 3.83 (br, 1?), 2.12 (m, 1H), 0.91 (d, *J* = 7.5, 3H), 0.81 (d, *J* = 7.5, 3H); ^13^C NMR (125 MHz, DMSO-*d*β) *δ*_C_ 165.5, 158.7, 134.6, 98.9, 60.4, 33.2, 18.0, 16.5.

Cyclic-l-Pro-l-*O*Met (A13) was dissolved in 50 μL of 30% MeOH/H_2_O solution. Cyclic-l-Val-ΔAla (A19) was solved in 70% MeOH/H_2_O solution. An aliquot (10 μL) of each DKP solution was added to 1 mL of L1 media containing *P. tricornutum* cells. Cell counts were taken at day 3 and 7. Three biological replicates were used for each cell count (Figure 3).

### Marfey’s method for determination of absolute configurations of 1 and 2

1 mg of cyclic-l-Pro-l-*O*Met (**1**) and cyclic-l-Val-ΔAla (**2**) were hydrolyzed in 0.5 mL of 6 N HCl at 115 °C. After 1 h, the reaction mixtures were cooled to rt by placing on ice water for 3 min. The reaction mixture was evaporated *in vacuo* then the material was resuspended in 0.5 mL H_2_O and dried under vacuum (x 3) to ensure complete removal of the acid. The hydrolysate of each DKP was lyophilized overnight. Subsequently, the dried reaction material and 0.5 mg of each amino acids, including l-Pro, d-Pro, l-*O*Met, l/d-*O*Met, l-Val and d-Val, were each dissolved in 100 μL of 1 N NaHCO_3_, followed by the addition of 50 μL of 10 mg/mL l-FDAA (1-fluoro-2,4-dinitrophenyl-5-l-alanine amide) in acetone. The reaction mixture was incubated at 80 °C for 3 min then quenched by the addition of 50 μL of 2 N HCl. Then 300 μL of 50% ACN/50% H_2_O was added to each mixture and 10 μL was analyzed by LCMS using a Kinetex® EVO C_18_ column (100 × 4.6 mm, 5 μm) using the following gradient solvent system: gradient from 20% ACN + 0.1% FA/80% H_2_O + 0.1% FA to 60% ACN + 0.1.% FA/40% H_2_O + 0.1% FA over 40 min with a flow rate of 0.7 mL/min (UV detection at 340 nm). The retention times of authentic acid l-FDAA derivatives l-Pro (13.1 min), d-Pro (16.8 min), l-*O*Met (9.7 min), d-*O*Met (10.6 min), l-Val (20.7 min) and d-Val (24.2 min); the hydrolysate products gave peaks with retention times of 13.1, 9.8, and 20.7 min, according to l-Pro, l-Val, and l-*O*Met, respectively.

### Total synthesis of cyclic-l-Pro-l-*O*Met (1)

Boc-l-methionine-sulfoxide (1000 mg, 3.76 mmol) was dissolved in dichloromethane (20 mL) followed by *O*-(Benzotriazol-1-yl)-.*N..N..N’..N*’-tetramethyluronium hexafluorophosphate (1.713 mg, 4.52 mmol) and the reaction mixture was stirred for 30 min at rt. Subsequently, l-Proline methyl ester (749 mg, 4.52 mmol) and triethylamine (0.63 mL, 4.52 mmol) were added, and the mixture was stirred for 18 h at 4 °C in a cold room. After 18 hours, the reaction mixture was quenched by addition water (10 mL). The aqueous layer was extracted with ethyl acetate (20 mL x 3). The organic layer was dried under vacuum to yield 580 mg of crude white oil (41% yield). The oil was then dissolved in 1 mL of anhydrous dioxane, heated to 50 °C and 3 N hydrochloride (0.6 mL, ~4 equiv.) was added dropwise. The reaction mixture was stirred for 3 h and then worked up by evaporating the solvent under vacuum. The mixture was then lyophilized under high vacuum overnight. The final cyclization reaction was accomplished by resuspending the dried oil in anhydrous dimethylformamide (2 mL) then heated 100 °C and stirred for 4 h. The mixture was frozen and then dried by lyophilizing under high vacuum overnight. The resulting oil was purified by reversed phase prep-HPLC using a Luna® Phenyl-Hexyl column (250 × 21.2 mm, 5 μm) with the following solvent gradient system: 5% ACN/95% H_2_O to 20% ACN/80% H_2_O over 40 min with a flowrate of 10 mL/min. Cyclic-l-Pro-l-*O*Met (**1**) was obtained as white oil with an overall yield of 26.6% (250 mg).

### Absolute quantifications of the two DKPs in different strains

Production levels of **1** and **2** produced by different *B. thuringiensis* strains – *B. thuringiensis israelensis*, *B. thuringiensis* sp. 4A4, *B. thuringiensis* sp. 4Q7, and *B. thuringiensis* sp. 407 – were quantified using 1 L cultures of each strain. 5 mL of overnight LB cultures of each strain were used to inoculated 500 mL of L1 media. Cultures were grown at 22 °C without shaking for 21 days. Then each supernatant was centrifuged (7000 rcf, 30 min) and freeze-dried. The dried cell pellets were weighed in order to normalize production levels to total biomass. The spent supernatant of each extract was passed over a column containing 20 g of XAD4 and XAD7 resin mixture the crude extract was eluted with 40 mL of DI water and 40 mL of 100% MeOH. The dried crude extract was then resuspended in approximately 20 mL of 100% DI water and was loaded on Sep-pak C_18_ cartridge (Waters Corporation). The cartridge was washed with 20 mL of 100% H_2_O and 20 mL of 25% MeOH/75% H_2_O and dried to form one fraction per strain. The fractions were resuspended in 500 μL of 50% MeOH/50% H_2_O and 10 μL was analyzed on the LCMS equipped with a Kinetex® EVO C_18_ column (100 × 4.6 mm, 5 μm) using the following gradient system: 5% ACN/95% H_2_O to 60% ACN/40% H_2_O over 20 min at a flow rate of 0.7 mL/min. Integrated extracted ion chromatograms (EIC) for ion adducts corresponding to **1** (*m*/*z* 244) and **2** (*m*/*z* 168) were analyzed and compared to a six-point standard curve generated by authentic standards.

### *P. tricornutum* growth stimulation by DKP and mixed single amino acids

Cyclic-l-Pro-l-*O*Met (**1**) was dissolved in 30% methanol in water at a stock concentration of 100 mM. cyclic-l-Val-ΔAla (**2**) was dissolved in 70% methanol in water at a stock concentration of 20 mM. 30% and 70% methanol solutions were used in DKP-free control cultures respectively. Final concentrations were obtained by diluting DKPs in the L1 media. *P. tricornutum* cells were cultured in 1 mL of L1 medium containing 10mM, 2mM, 1mM, 100μM, 10μM, 1μM and 100nM final concentrations for **1**. 700μM, 350μM, 35μM, 3.5μM, 350nM and 35nM final concentrations for **2**, and a DKP-free methanol, respectively. Counts were taken on day 7. Solutions containing mixed pair of single l-amino acids (l-Pro/l-Met AA and l-Val/l-Ala AA) were added to L1 media. l-Pro/l-Met AA were added concentrations of 10mM, 1mM, 100μM and 10μM, l-Val/l-Ala AA was added at concentrations 700μM, 350μM, 35μM and 3.5μM. Counts were taken on day 8.

### RNA-extraction

10 mL culture consisting of 2.6 E+7 of *P. tricornutum* cells were added at Day 0 in 50 mL Falcon tubes with a final concentration of 100 μM cyclic-l-Pro-l-*O*Met, or a DKP free control (10 μL 30%/70% Methanol/H_2_O). Cells were harvested on days 3 and 6 as 10 mL cultures. Cells were lysed by sonication using the Microson Ultrasonic Cell Disruptor (Misonix) at setting 12 for 10 x 1-sec pulses. RNA was extracted from the cells using a CTAB method (Hawkins et al., 2016).

### Bioinformatic analysis

#### RNA-Seq data preprocessing

A total of 12 samples (4 conditions x 3 biological replicates) were sequenced, each with two FastQ files (paired-end of 150 bp), generated a total of 24 RNA-seq files. Each file ranges from 34.9 to 49.0 million reads. The quality of the RNA-Seq data was examined by Fastqc (v0.11.8) (Andrews, 2010). Multiqc (v1.7) (Ewels et al., 2016) summarized the reports generated by Fastqc. Trimmomatic (v0.39) trimmed the adaptors and low-qualify bases. Reads shorter than 36 were discarded.

#### Gene Quantification

The isoform expression level was estimated by Salmon (v0.11.2). The reference transcriptome was downloaded from Ensembl (Yates et al., 2020). Salmon index was built using k-mer size 31, and duplicated sequences in the reference transcriptome were kept. The RNA-Seq reads were quantified against the index using Salmon mapping-based mode. --seqBias flag was passed to Salmon for correcting sequence-specific biases. Tximport (v1.10.1) (Soneson et al., 2016) summarized the isoform abundance for gene-level analysis.

#### Differential expression analysis

Differentially expressed genes between different conditions were detected by DESeq2 (v1.22.2) (Love et al., 2014) using 11 of the 12 samples (one sample deviates from the other two replicates (Figure S2) and was not used in the analysis). Genes with low reads counts were filtered. The genes with *padj* < 0.05 and *|log_2_FoldChange*| > 1 were considered as differentially expressed genes. The principle component analysis was performed by the built-in function plotPCA in DESeq2. The expression patterns of the differentially expressed genes were depicted by heatmaps that were generated using R package pheatmap (v1.0.12) (https://cran.r-project.org/web/packages/pheatmap/index.html) The expression value was represented by *log_2_(TPM* + 1).

#### GO enrichment analysis

The GO terms associated with genes were retrieved by the Retrieve/ID mapping tool in UniProt (2019). TopGO (v2.34.0) (Alexa and Rahnenfuhrer, 2019) was applied to identify the enriched GO terms. Fisher’s exact test was used in TopGO to determine whether the assigned GO categories are significant. The p-value cutoff was 0.05.

#### Co-expression network analysis

WGCNA (v1.66) (Langfelder and Horvath, 2008) and consensus clustering approach (Monti et al., 2003) were combined to build robust consensus co-expression networks (Shahan et al., 2018). WGCNA was ran 1000 times with randomized parameters and subsampled genes. Genes with low variance and zero MAD were filtered. The expression level was measured as *log_2_ (TPM* + 1). Biweight midcorrelation was selected as the correlation method. And signed network was chosen as the network type. A consensus matrix was generated by dividing the number of times genes were clustered together by the number of times genes were subsampled together. The final co-expression network was constructed from the consensus matrix by running WGCNA with power 6 and minModuleSize 10. The module eigengenes were calculated by WGCNA and presented by heatmap. And the top 5 GO terms enriched for the interesting gene modules were shown by bar graphs.

### Fatty Acid analysis and BODIPY staining

2.5mL of mid-exponential *P. tricornutum* cells was added to 50 mL of L1 medium or 50 mL of L1 medium containing *B. thuringiensis* lysate (see earlier section on *B. thuringiensis* lysate production). After 7 days, cells were centrifuged 13,300 rpm for 30 seconds, and the pellet was frozen in liquid nitrogen and stored at −80 °C. To extract lipids, a mixture of 628 μL methanol, 251 μL chloroform, and 251 μL water was added to the pellet. This suspension was vortexed and held at −20 °C for 90 min, with further vortex at 30 and 60 min. Extracts were centrifuged at 8,000 rpm for 5 min and the supernatant was transferred to a separate vial. An additional mixture of 286 μL methanol and 286 μL chloroform was added to the cell pellet. The sample was vortex mixed and held at −20 °C for 30 min, centrifuged at 8,000 rpm for 5 min, and the supernatant was added to the previously saved supernatant. 286 μL of water was added to the combined liquid extract to achieve phase separation. The aqueous methanol layer was removed from the chloroform. The chloroform was evaporated under a gentile stream of nitrogen, and lipids were derivatized by adding 200 μL of 3 N methanolic HCl to the dried chloroform layer and heated at 70 °C for 1 h. After allowing the sample to cool to room temperature, 100 μL of hexanes were added and the sample vortex mixed. The upper hexane layer containing the methyl-esterified lipids was collected. An additional 100 μL of hexanes was added to the acid raffinate and vortex mixed. The upper hexane layer was again collected and combined with the first extract. 50 μg methyl heptadecanoate was added as an internal standard. All samples were analyzed using a Bruker 450-GC gas chromatograph equipped with a Varian VF-5ms (30 m × 0.25 mm × 0.25 μm) column coupled to a Bruker 300MS mass spectrometer (Bruker, Fremont, CA) in full scan, electron ionization mode. The GC oven temperature was initially set at 150 °C for 2 min, raised to 205 °C at 10 °C min^-1^, then raised to 230 °C at 3 °C min^-1^, and finally raised to 300 °C and held for 3 min at 10 °C min^-1^. Compounds were identified using the Varian MS workstation software (version 6.9.3) in conjunction with the NIST mass spectral library (National Institute of Standards and Technology, Gaithersburg, MD). Ion intensities were quantified using the sum of the total ion chromatograph peak area. The signal intensity of each compound within a sample was first normalized to the intensity of the internal standard, and then normalized to the total cell number in each sample.

For BODIPY 505/515 (ThermoFisher) staining of lipids, *P. tricornutum* was grown in L1 medium, L1 + *B. thuringiensis* lysate, or L1 + 100 μM cyclic-l-Pro-l-*O*Met (**1**). 1 mL of each culture was harvested at 12 days (stationary phase) and placed in a microcentrifuge tube. 1μl of 10 mM BODIPY (dissolved in DMSO) was added to each tube, which was left in the dark for 10 minutes. Afterwards, *P. tricornutum* cells were washed twice with fresh L1 medium (pelleted at 11,000 g for 10 min.) and then imaged with a Leica SP5X confocal microscope. The confocal images were analyzed using ImageJ-Fiji (https://imagej.net/Fiji). To quantify fluorescence, acceptable fluorescence intensity threshold was set between 50 and 225. A low threshold of 50 removes background fluorescence, while the high threshold of 225 excludes potentially oversaturated pixels. Individual cells were circled; Integrated Density (Mean Fluorescence Intensity x Area of Fluorescence with readings between 50 and 225) was obtained for the circled cell and plotted in Figure 6b. DIC and fluorescent images of individual *P. tricornutum* cells shown in Figure 6a were obtained at 250x magnification using a Zeiss Axioscope with a Sony Alpha 7mrII camera (2.5x).

## Acknowledgement

We are grateful for advice, bacterial strains, microscopy, equipment provided by Drs. Charles Delwiche, Eric Haag, Jonathan Goodson, Lei Guo, Cordelia Weiss, Caren Chang, Vince Lee, and Steven Farber. We would like to thank the dedicated assistance by undergraduates Ms. Caroline Gonter, Marta Roman, and Lina Sobh. This work has been supported by a grant from NSF/CBET 1134115 to G. S. and Z. L. as well as the USDA National Institute of Food and Agriculture, Hatch project 1010278 to Z. L. J. S. was supported by the Wayne T. and Mary T. Hockmeyer Endowed Fellowship. M. L. was supported in part by NSF award DGE-1632976.

## Author Contributions

J.S., M.B., E.M., A.Q., performed the experiments. M.L. performed the RNA-seq analyses. J.S., M.B., E.M., and Z.L. wrote the manuscript. J.C., G.S., and Z.L. guided the study. Z.L. agrees to serve as the author responsible for contact and ensures communication. All authors read, revised, and approved the manuscript.

